# Maskless and on-chip LED-array microscope with spatially-varying angle calibration for centimeter-scale phase imaging

**DOI:** 10.1101/2025.09.05.674268

**Authors:** Sibi Chakravarthy Shanmugavel, Vindya Senanayake, Donghwa Suh, Jeffrey M. Gross, Shwetadwip Chowdhury

**Affiliations:** Chandra Department of Electrical and Computer Engineering, University of Texas at Austin, Austin, TX, USA; Department of Molecular Biosciences, University of Texas at Austin, Austin, TX, USA

## Abstract

On-chip imaging with LED-array-based angled illumination offers a cost-effective approach for large field-of-view (FOV) phase imaging. However, it faces two main challenges: (1) twin-image ambiguity can degrade phase reconstruction. While mask-based modulation can help, it adds system complexity due to fabrication and alignment requirements; and (2) the illumination angle from each LED varies across large FOVs, and can degrade centimeter-scale phase reconstruction without calibration. Here, we present a computational framework to jointly achieve mask-free on-chip phase imaging and adaptive calibration of spatially-varying illumination angles. The sensor FOV is divided into subregions within each of which LED illumination is approximated as planar. LED illumination angles for each subregion are initialized geometrically. Phase retrieval is then performed within each subregion by constraining the reconstruction with a soft optical transparency prior while simultaneously refining angle estimates. Reconstructed phase maps are merged to produce a high-quality, large-FOV phase image. We demonstrate this approach by achieving centimeter-scale on-chip phase imaging (up to 2.7 × 1.7 cm^2^) with micron-level resolution across various biological tissue sections.

## 1. Introduction

Optical microscopes are essential tools in biological sciences, supporting applications like disease diagnosis, cellular studies, and tissue engineering [1-3]. However, conventional systems are constrained by a fundamental trade-off between resolution and field-of-view (FOV), which limits overall imaging throughput [4]. This trade-off is often characterized by space-bandwidth product (SBP), which is a measure of the imaging system’s information-carrying capacity, as defined by the number of independent resolvable points in the image based on both the system’s resolution and FOV. In practice, standard microscope objective lenses offer SBPs ranging from a few megapixels at high numerical aperture (NA) (i.e., resolution) to several tens of megapixels at lower NA [5]. This limitation typically forces one to compromise between visualizing a sample at high resolutions or over large FOVs, and presents a major challenge for applications where important submicron features span across millimeter-to centimeter-scale regions. A common solution is to mechanically scan the sample through the small FOV of a high-NA objective using motorized stages; however, this method can be slow, expensive, and prone to alignment errors and optical aberrations.

Recent advances in computational imaging have enabled gigapixel-scale imaging by surpassing the traditional limits of SBP. For example, Fourier ptychography [6, 7] and structured illumination microscopy [8-10] leverage angled and/or patterned illumination to encode raw measurements with high-resolution sample information that is traditionally inaccessible by the system’s limited NA. By decoding this information, these methods computationally improve resolution across the entire FOV, effectively increasing the imaging SBP. Another approach involves using multiple cameras, each equipped with an individual lens, to simultaneously capture adjacent regions of the sample [11-13]. With proper calibration and alignment, the resulting image data can be seamlessly stitched together to form a continuous and wide FOV. Additionally, overlapping regions between neighboring views can be leveraged to generate stereoscopic reconstructions, enabling topographical visualization of the sample [12].

However, the high-SBP computational imaging techniques mentioned above still rely on lenses, and therefore face challenges associated with bulky and costly system design. Moreover, achieving a large FOV is best achieved with low-to moderate-NA lenses, which compromises axial resolution and thus poses a significant challenge for high-resolution 3D microscopy applications. A separate class of computational imaging methods has emerged that addresses these challenges by removing the optical lens altogether [14, 15]. These lensless imaging systems often operate in a simple and compact “on-chip” configuration, where the sample is directly placed on a 2D sensor. In this setup, a small gap of < 1 mm usually remains between the sample and the active sensor, and the imaging FOV is directly dictated by the sensor’s physical size (typically centimeter-scale). Although spatial resolution is constrained by sensor’s pixel size (typically microns), pixel super-resolution techniques [16-21] can overcome this limitation, enabling subcellular-resolution imaging across centimeter-scale FOVs.

Lensless on-chip imaging systems can also support multiple modes of contrast, such as phase [22, 23] and fluorescence [24-26]. Here, we specifically focus on quantitative phase imaging (QPI), which enables quantitative label-free imaging of morphological features in transparent specimens (e.g., cell cultures, thin tissue sections, etc). By combining high-SBP, label-free sensitivity, and a compact system design, on-chip QPI systems offer a powerful platform for a wide range of biological applications, such as cell counting [27, 28], microfluidics [29], histology [30], pathology studies [31], and air quality monitoring [32]. Notably, because image sensors are inherently sensitive to only light intensity, recovering the phase of the light transmitted through the sample requires computational phase retrieval. This process typically involves capturing multiple measurements across various perturbed imaging configurations, which introduce diverse sample-specific information into the measurements. With sufficient measurement diversity, computational algorithms can accurately reconstruct a complex-valued amplitude and phase image of the sample simply from raw intensity measurements.

To collect diverse measurements with lensless on-chip systems, various strategies are employed. For example, capturing measurements at varying sample-to-sensor distances leverages optical defocus as the primary source of measurement diversity [22, 33, 34]. Separately, ptychographic techniques capture multiple measurements with a laterally translating mask or diffuser element, encoding measurement diversity through modulated illumination patterns to the sample [28, 35, 36]. Both optical defocus and ptychography-based approaches often use motorized mechanical stages with high precision and repeatability, making them expensive and susceptible to reconstruction artifacts caused by specimen drift and mechanical errors. Pixel-programmable devices, such as spatial light modulators [37, 38] or digital micromirror devices [39], can be used to modulate illumination patterns without mechanically moving the sample. However, such systems are typically expensive and require lenses to project the patterns onto the sample. This compromises the simplicity and compactness of the setup. Other methods have emerged that capture images at different wavelengths [17, 18, 40-42]. These multi-wavelength datasets can then be used to recover a sample’s phase information, also without any mechanical movement. However, this strategy relies on the sample’s refractive-index remaining constant across wavelengths (i.e., no dispersion). In practice, refractive-index is known to be dependent on wavelength, and thus additional wavelength calibration and dispersion compensation may be required to achieve truly quantitative results [17, 42].

Measurement diversity in lensless on-chip systems can also be achieved by capturing sample measurements at various illumination angles. A practical and cost-effective way to accomplish this without mechanical movement, physical lenses, or expensive wavelength tunable lasers is to use an LED array. By activating individual LED elements in the array, the sample is naturally illuminated at different angles [7, 41, 43, 44]. However, though the hardware is simple and scalable, phase retrieval remains a challenge due to the problem’s underdetermined nature. Specifically, phase retrieval seeks to reconstruct the sample’s complex valued image (i.e., amplitude and phase) from raw intensity (i.e., amplitude squared) measurements captured by the sensor, which discard crucial phase information. This loss of information can lead directly to the twin-image problem, which often results in ambiguity and stagnation in the phase retrieval reconstruction process [45-47]. To mitigate this, additional constraints are typically incorporated into the optimization process. One common method is the synthetic aperture constraint in the Fourier domain [30], analogous to the pupil function constraint used in Fourier ptychography [7], where reconstruction relies on spectrum overlap between adjacent measurements. However, this approach is inherently two-dimensional and thus limited in its ability to reconstruct three-dimensional samples. As mentioned above, another effective method is to use a mask, positioned adjacent to the sample, to controllably modulate light before it reaches the sensor [47]. In combination with an LED array, light modulations from the mask can be translated based on the angle of illumination, thus imposing an additional spatial constraint that improves the posedness of the phase retrieval process without any mechanical movement. Building on this concept, later studies have explored modulation strategies using amplitude masks [48, 49], phase masks [44], pinholes [50], blood-coating [35, 36], and diffusers [41, 51] to further enhance robustness and reconstruction quality.

However, mask-based modulation introduces notable challenges: (1) characterizing a mask’s modulation pattern can be time-consuming; (2) fabricating custom masks, particularly at tight tolerances, can be expensive; (3) masks often reduce light throughput, leading to longer exposure times; and (4) complex mask structures can be difficult to align precisely. This alignment is especially critical in nonconvex computational imaging frameworks, where even minor misalignments (e.g., environmental drift) can significantly degrade reconstruction quality and necessitate frequent recalibration. These limitations compromise the key simplicity, compactness, and affordability benefits of LED array-based on-chip systems.

Another often-overlooked challenge is with the angular illumination provided by a LED array. Each LED in the array is commonly assumed to provide uniform, plane-wave illumination at a fixed angle across the entire sample [30, 41, 44, 48-51]. In reality, LEDs more closely emit spherical waves, producing near-normal incidence at the center of the beam and increasingly oblique angles towards the edges. As a result, a single LED illuminates different regions of the sample at varying angles. This effect is further compounded by any residual tilt or rotation of the array relative to the sensor. Though negligible for small FOVs, this effect can introduce significant artifacts across the large FOVs required for high-SBP imaging. To the best of our knowledge, this challenge has not yet been addressed in lensless on-chip imaging systems.

In this work, we present a novel phase-retrieval framework for on-chip phase imaging using multi-angle illumination from an LED array. *Importantly, our approach eliminates the need for mask-based modulation and explicitly accounts for the spatial variation in illumination angle that occurs due to each LED illuminating centimeter-scale regions in the sample*. We validate our framework by reconstructing gigapixel-scale phase images (up to > 2 gigapixels, spanning 2.7 × 1.7 cm^2^ with micron-level resolution) of biological specimens that exhibit densely organized structures with rich phase contrast, such as adult mouse kidney, zebrafish head sections, and whole mouse embryos. All measurements were acquired with a simple board-level sensor illuminated by a commercially available LED array.

## 2. Methods: Optical system and pre-processing

We first describe our physical on-chip phase-imaging system, as well as the pre-processing steps taken to align and characterize the system before conducting joint on-chip phase-retrieval and angle calibration.

### A. Optical setup

A schematic of our on-chip LED array microscope is illustrated in **Figure 1(a)** below. The setup features a programmable LED array (Adafruit Inc.) with a 32 × 32 grid of independently controllable LEDs, each spaced 4 mm apart with an approximate element size of ∼120 µm. The LEDs provide illumination at three wavelengths: 632 nm (red), 520 nm (green), and 467 nm (blue), each with a 40 nm bandwidth. The matrix is positioned approximately 30 cm (h≈30 cm) above the sample, which is placed near the sensor at a distance of approximately 1mm.

**Figure 1:**
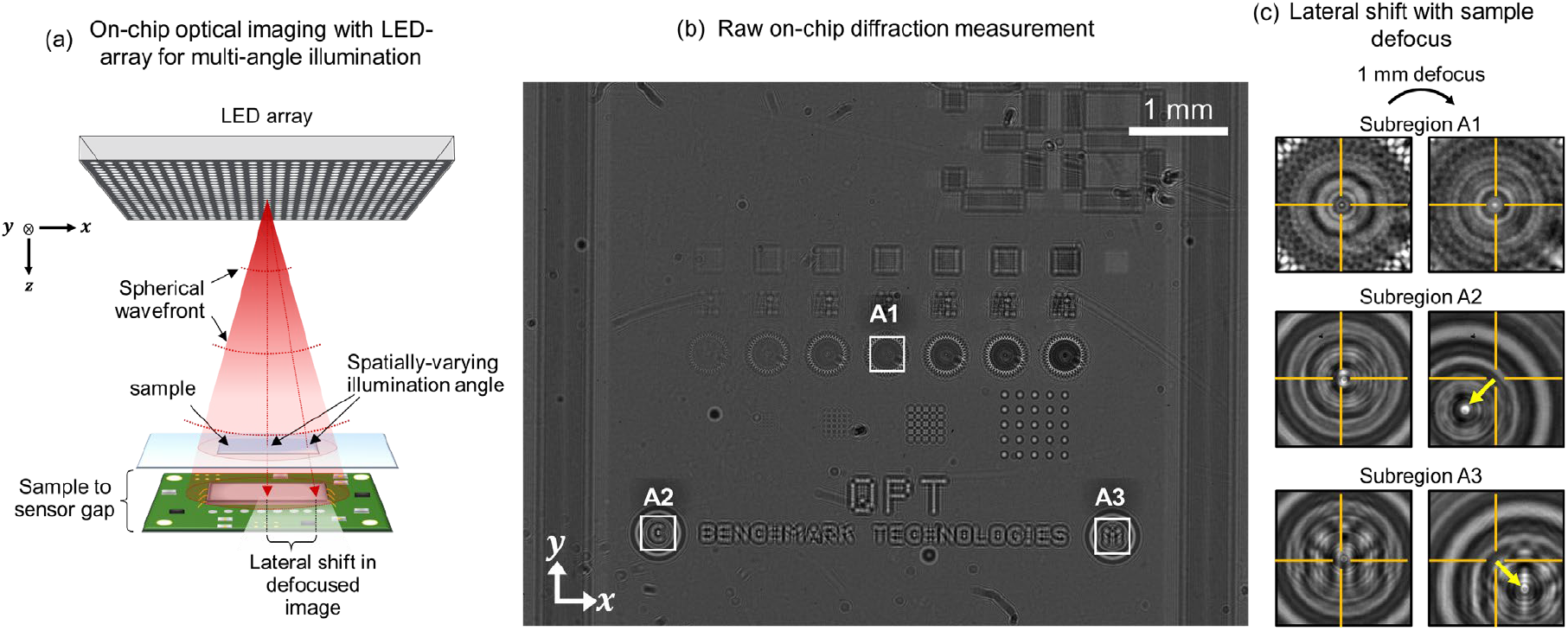
**(a)** Simplified schematic of our lensless on-chip imaging system with multi-angle illumination. A programmable LED array, positioned ∼30 cm above the sample, provides quasi-monochromatic illumination at three wavelengths (632 nm, 520 nm, and 467 nm). A bandpass filter (not shown) is placed above the sample to improve temporal coherence at 632 nm, the wavelength used in our experiments. A board-level monochrome camera captures the resulting diffraction patterns. The LED array is controlled by a Raspberry Pi, which also synchronizes LED activation with image acquisition via a trigger pulse. **(b)** Full FOV diffraction pattern of a phase target illuminated by a reference LED. A Siemens star pattern at the center (subregion **A1**) is used as the reference subregion for alignment. **(c)** Diffraction patterns from subregion **A1** before and after 1 mm axial defocus exhibit no lateral shift, confirming orthogonal (on-axis) illumination. In contrast, subregions **A2** and **A3** located farther from the center of the LED’s axis receive increasingly angled illumination and exhibit noticeable lateral shifts in their diffraction patterns upon defocus. This lateral displacement reflects the spatial variation in illumination angle, which becomes more pronounced with increasing distance from the reference LED’s optical axis.

This configuration enables the LED illumination to achieve sufficient spatial coherence. For image acquisition, we use a monochrome board-level camera (DMM 37UX226-ML, The Imaging Source Inc.) with a 4000×3000 pixel resolution and 1.85 µm pixel size, yielding a 7.4 mm × 5.55 mm FOV across the sensor. The LED array is controlled by a Raspberry Pi microcontroller equipped with an RGB matrix bonnet. Camera synchronization with the LED array is achieved through a trigger pulse generated by the Raspberry Pi during each LED activation. To achieve quasi-monochromatic illumination with sufficient temporal coherence, a 10 nm full-width at half-maximum (FWHM) bandpass filter (Edmund Optics # 65-227), centered around the chosen illumination wavelength, is placed directly above the specimen.

### B. LED array and sensor alignment

To streamline the angle calibration process, we found it effective to: 1) divide the total measurement FOV into subregions, each with size smaller than the coherence area of the illumination (see **Supplementary Note S1.1**); and 2) designate a ‘reference’ subregion at the center of the sensor’s FOV, which we align on-axis with a single ‘reference’ LED at the center of the LED array. This ensures that the reference subregion receives essentially on-axis plane-wave illumination from the reference LED (see **Supplementary Note S1.2**). This will be useful for later estimating sample-to-sensor distance and initializing illumination angles for other remaining LEDs in the array. This alignment step is straightforward and is most easily performed by imaging a symmetric 2D object at two or more defocus planes. If properly aligned on-axis to the LED, defocusing should not cause any lateral shift in the center of the captured diffraction pattern. We demonstrate this in **Figure 1** using a Siemens star pattern from a phase calibration target (Benchmark Technologies). The Siemens star, located in subregion A1 at the center of the FOV [**Figure 1(b)**], was selected as the alignment reference due to its radial symmetry. The LED array was then laterally adjusted until axial defocus no longer produced lateral displacement in the diffraction pattern of the Siemens star [**Figure 1(c)**]. This confirmed that subregion **A1** was receiving orthogonal illumination from the reference LED, indicating successful optical alignment.

### C. Determining distance between sample and sensor

Using this on-axis illumination in the reference subregion, we next determine the gap distance between the sample and the sensor. We leverage the property that thin, transparent samples, such as phase targets or tissue sections, can be modeled as 2D phase-only objects. At the plane of focus, these samples do not induce amplitude modulation; such fluctuations arise only due to defocus. Since there is a gap between the sample and the sensor, the measured diffraction patterns inherently contain amplitude variations caused by this defocus. To estimate the gap distance, we computationally propagate the measured amplitude within the reference subregion (illuminated by the reference LED) across a range of distances and identify the plane where the amplitude modulation is minimized, corresponding to the true object plane. This is achieved by finding the propagation distance that minimizes the Brenner gradient of the amplitude pattern [52] (see **Supplementary Note S1.3**).

### D. Determining global LED coordinates

We next estimate the global transverse coordinates of each LED in the array relative to the reference LED, which we defined earlier as the origin. A key observation is that small changes in illumination angle produce lateral shifts in defocused diffraction patterns that are directly related to the magnitude and direction to the angle change itself. To determine each LED’s position, we sequentially activate off-axis LEDs and measure the resulting lateral shifts via cross-correlation. This is done within the central reference subregion, which is known to receive on-axis plane-wave illumination from the reference LED. By comparing each off-axis measurement to the one obtained from the reference LED, and using the previously computed sample-to-sensor distance, we compute the global spatial coordinates of each LED using simple geometric relationships (see **Supplementary Note S1.4**).

### E. Geometric initialization of spatially-varying illumination angles for each LED

Finally, we compute the illumination angle that each LED imparts to each subregion of the sample. This spatial variation arises because LEDs emit spherical, not planar, wavefronts. However, since each subregion is smaller than the illumination’s coherence area, the light incident to that subregion can be locally approximated as a plane wave. Subregions in the sample located farther from an LED’s optical axis experience increasingly oblique plane-wave illumination from that source [subregions **A2** and **A3** of **Figure 1(b, c)**]. With the global LED coordinates already determined, the illumination angle from any LED to any subregion can be efficiently estimated using simple geometric calculations based on the direct path connecting the LED to the subregion center (see **Supplementary Notes S1.5**). These estimates are then later refined through joint variable optimization, as described below.

## 3. Methods: Computational framework

We now describe our computational joint-variable optimization framework for on-chip phase retrieval and adaptive calibration of illumination angles. To do this, we present a computational forward model 𝒢that predicts on-chip diffraction measurements from a sample’s 2D complex transmittance, incorporating key system parameters such as illumination angle, sample-to-sensor distance, and sensor pixel size. Notably, due to spatial variations in the illumination angle across the sensor’s large FOV, the model is applied independently to each subregion of the sample, within which the illumination can be approximated as a plane wave with constant angle. Thus, the forward model predicting the scattered measurement at the *n*-th subregion under illumination from the *𝓁*-th LED can be expressed as:

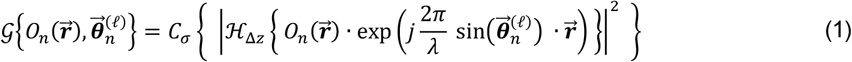

Here, 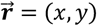 denotes the 2D spatial coordinate vector and 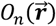 represents the 2D complex transmittance of the sample’s *n*-th subregion. 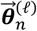 is the 2D vector describing the angle of incidence from the *𝓁*-th LED onto the *n*-th subregion. ℋ_Δ*z*_{·}denotes field propagation over the sample-to-sensor distance Δ*z*, implemented using angular propagation [53]. A detailed description of this forward model is provided in **Supplementary Note S2**.

Computationally, 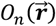 is a discretized map of the sample’s complex transmittance, upsampled by a factor *σ* and zero-padded to account for the maximum lateral shift induced by the largest illumination angle. Heuristically, *σ* is selected to approximate the ratio between the diffraction-limited resolution and the sensor’s pixel pitch. The operator *C*_*σ*_ {·}crops and downsamples (i.e., pixel-binning) the field derived from the optical operations on 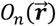, so that the dimensions of the forward model output match those of the raw on-chip measurements.

In practice, our goal is to recover the sample’s 2D complex transmittance from on-chip measurements. Additionally, we seek to adaptively refine the illumination angle estimates for each subregion, correcting for potential inaccuracies introduced by the earlier geometric method, which assumed purely lateral shifts under defocus as well as perfectly known system parameters. To achieve this, we formulate phase retrieval as a joint-variable optimization problem, which simultaneously refines estimates of the sample’s complex transmittance and the illumination angles. This is done by minimizing the mean squared error and the translational shift between model predictions and experimental measurements:

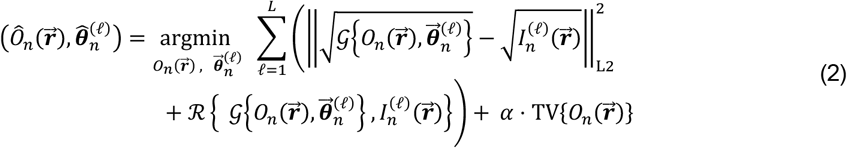

Here, 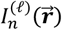 denotes the raw intensity measurement of the defocused pattern stemming from the *n*-th subregion being illuminated by the *𝓁*-th LED. The operator ℛ represents a correlation-based registration operation that outputs the translational shift between the model prediction and experimental measurement [54]. Total variation (TV) regularization is also implemented as a gradient sparsity-promoting prior for the sample. The parameter *α* denotes the strength of this regularization, and *L* represents the total number of LED elements.

At the start of the iterative reconstruction, the sample transmittance is initialized to a constant value of 1, corresponding to zero absorption and phase, while illumination angles are set to the geometric estimates computed earlier. To promote stable convergence without requiring mask modulation, we impose two major constraints:

1. Thin biological tissues are largely transparent with minimal absorption. We exploit this by applying a flat-amplitude constraint during early iterations, which narrows the solution space and improves stability. This constraint is lifted after a fixed number of iterations, allowing the amplitude to evolve freely for accurate recovery of the sample’s amplitude and phase.
2. Estimates of the illumination angle are iteratively refined using outputs from the registration operator ℛ. These updates are capped to a small pixel threshold (∼0.2 pixel) along the direction of the computed lateral shift. This prevents unstable behavior in early iterations. As the reconstruction progresses, the shifts naturally become small and the cap becomes unimportant, indicating stabilized reconstruction and refinement of angle estimates. This strategy, illustrated in **Figure 2** below, is implemented independently on each subregion of the sample. Once the iterative process converges, the recovered phase distributions from all subregions are merged together to generate the final high-SBP phase image.

**Figure 2:**
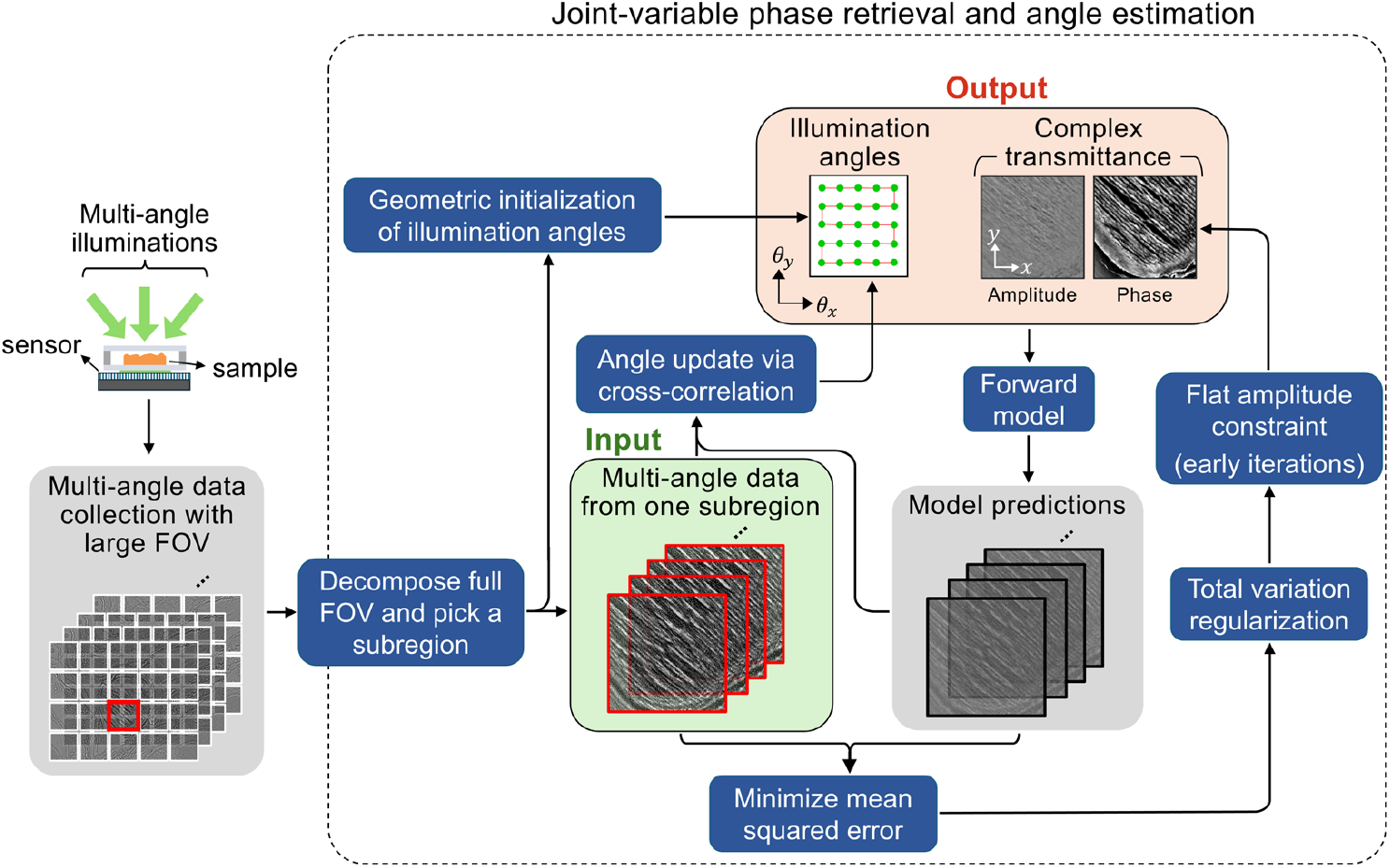
Workflow for joint-variable phase retrieval and angle estimation is illustrated for a single sample subregion. Raw intensity images are captured on-chip under multiple illumination angles. These multi-angle measurements are divided into subregions, each processed independently. With the multi-angle dataset from a single subregion, illumination angles are first estimated geometrically. A forward model then predicts the measured intensities based on the current estimates of both the angles and the sample’s complex transmittance. The mean squared error between predictions and measurements is iteratively minimized by incorporating gradient-based updates to the sample transmittance, angle refinements via cross-correlation, and application of a flat-amplitude constraint to the transmittance during early iterations. This cycle repeats for each subregion until convergence

A detailed description of the gradient derivation, step-by-step reconstruction procedure, as well as the compute resources used for reconstruction are provided in **Supplementary Notes S3, S4**, and **S5**, respectively.

## 4. Results

### A. Characterizing quantitative accuracy and imaging resolution

To assess the quantitative accuracy of our phase reconstructions, we imaged *D* = 10 μm diameter polystyrene microspheres. The microspheres had refractive index of *n*_PS_ = 1.5854 at our imaging wavelength of *λ* = 632 nm. The microspheres were suspended in index-matching oil with refractive index of *n*_*m*_ = 1.5684. The theoretical phase curve takes the form of a half-ellipse with diameter *D* = 10 μm, with peak phase value (2*π*/*λ*) · *D* · (*n*_pS_ − *n*_*m*_) = 1.69 radians.

**Figure 3(a,b)** show the reconstructed complex amplitude and phase reconstructions using a conventional optimization-based phase-retrieval algorithm [47, 55, 56] that uses stochastic gradient descent to estimate complex transmittance without additional constraints. As highlighted by the yellow arrows, this method results in severe artifacts and inaccurate phase reconstructions. The line plot in **Figure 3(e)** further confirms these limitations, with the reconstructed phase using conventional phase-retrieval diverging significantly from the ground truth. These findings are consistent with prior reports [44] and underscore the limitations of conventional methods, particularly their inability to robustly recover accurate phase without mask modulation.

**Figure 3:**
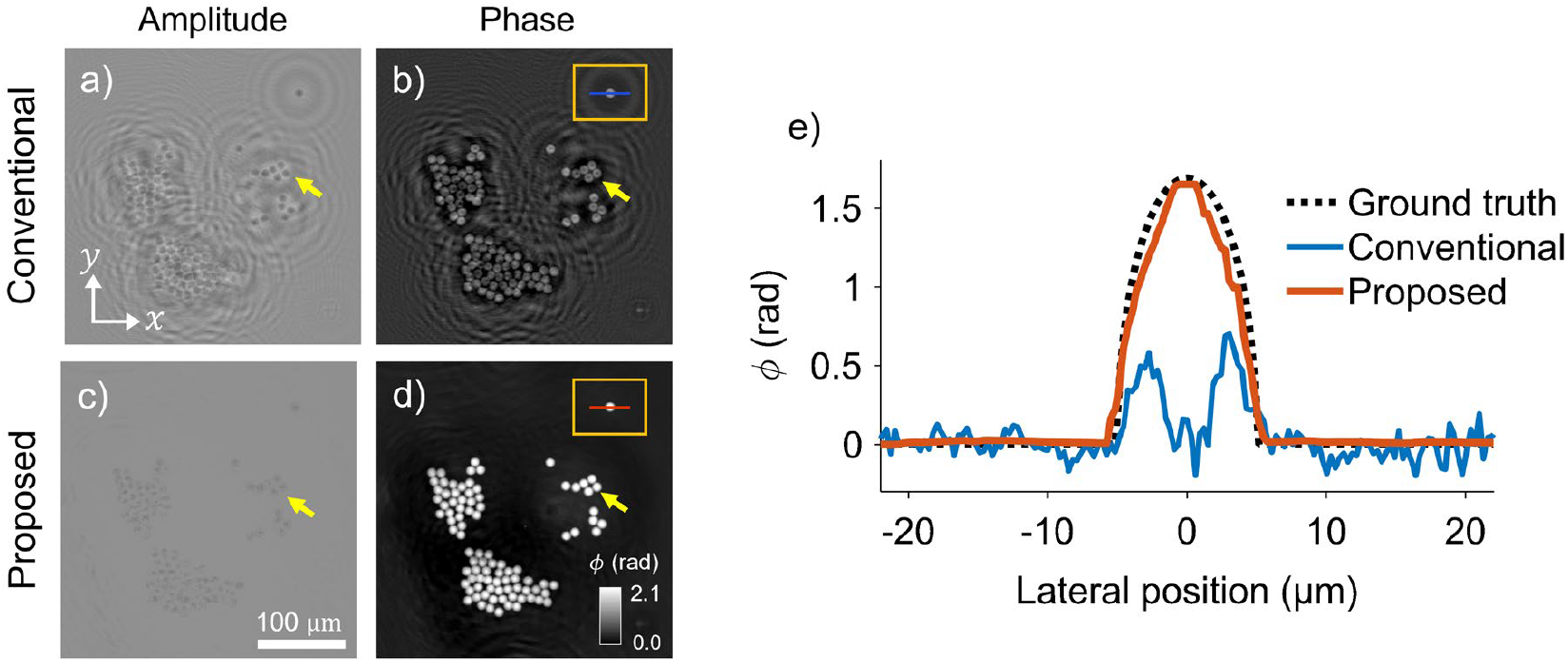
Quantitative phase reconstruction of 10 µm polystyrene microspheres. **(a, b)** Amplitude and phase reconstructions using a conventional on-chip phase-imaging method, showing artifacts and inaccuracies (yellow arrows). **(c, d)** Proposed method demonstrates improved amplitude and phase reconstruction, with accurate results highlighted by yellow arrows. **(e)** Line plot comparing the phase profile reconstructed with the conventional and proposed method. Conventional method shows significant mismatch with ground truth, while the proposed method closely matches the ground truth with < 2% error.

In contrast, our proposed method based on a regularized inversion with a flat-amplitude constraint yields substantially improved results, as seen in **Figure 3(c, d)**. The same regions highlighted by yellow arrows now exhibit clear and accurate reconstructions. The phase profile using the proposed method shown in **Figure 3(e)** closely matches the ground truth, with a maximum reconstructed phase delay of 1.66 radians, reflecting less than 2% error relative to the theoretical value of 1.69 radians. These results highlight the enhanced quantitative accuracy and robustness of our proposed lensless imaging approach.

### B. Characterizing imaging resolution

To assess our system’s imaging resolution, we imaged a USAF phase target (Benchmark Technologies). **Figures 4(a,b)** show the raw intensity measurement and the reconstructed phase image, respectively. Our on-chip system’s imaging resolution is governed by the effective numerical aperture (NA), which is calculated for using the following formula (see **Supplementary Note S1.6**):

**Figure 4:**
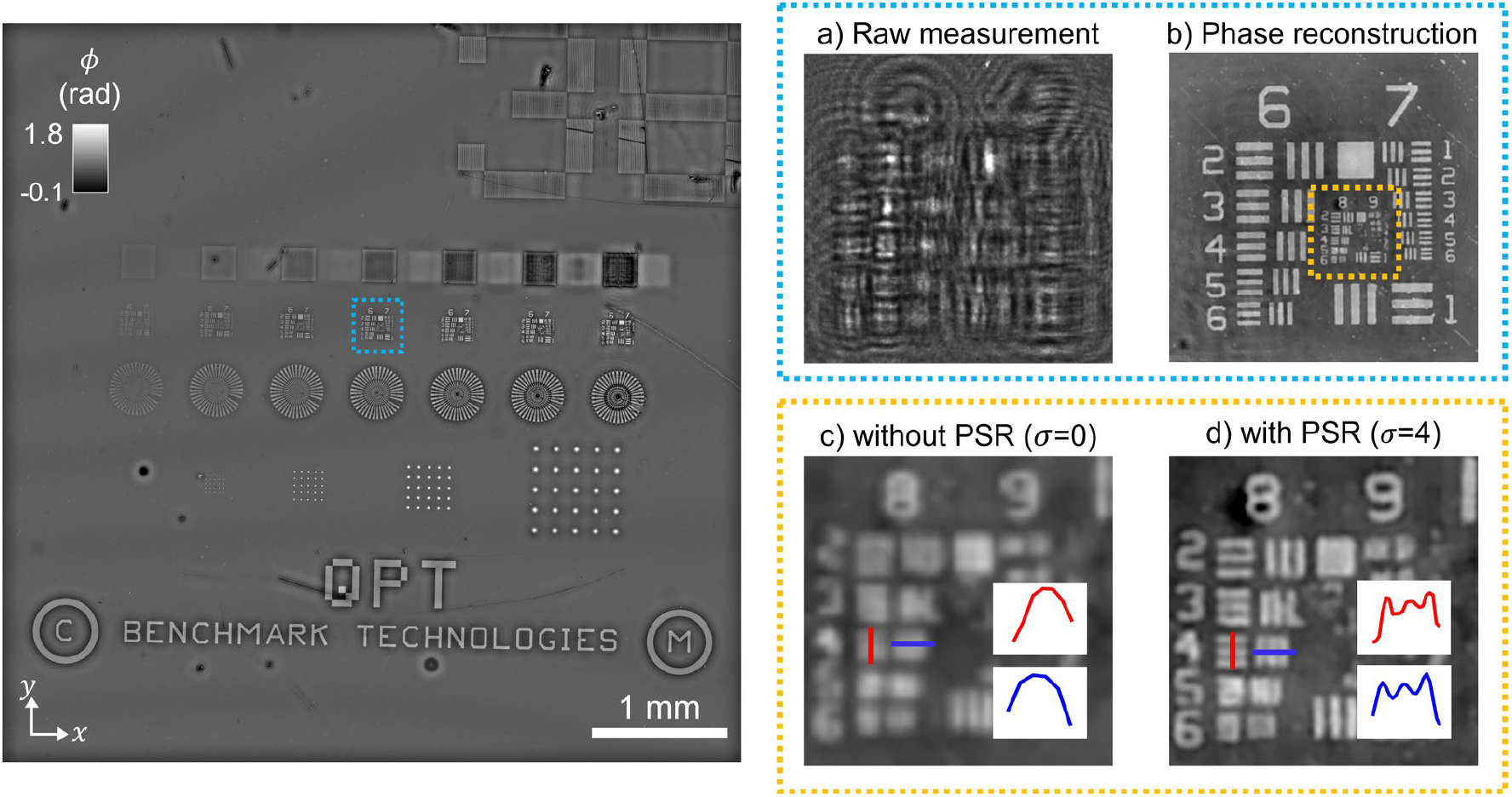
Resolution characterization using a USAF section of the total phase target. **(a)** Zoomed-in view from the dashed blue subregion of the full raw sensor measurement. **(b)** Reconstructed phase image showing clear resolution of fine features. **(c, d)** Further zoomed-in view, from the dashed yellow subregion in (b), shows image comparison without and with pixel super-resolution (PSR), respectively. Group 8 elements down to 1.55 µm are resolved only with PSR (*σ* = 4), demonstrating enhanced resolution beyond the sensor pixel limit.

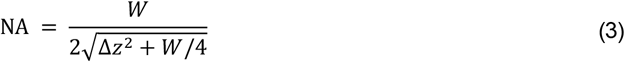

Recall that Δ*zz* denotes the sample-to-sensor distance, and *W* is the sensor width. For our system, these parameters yield a NA of 0.94, corresponding to a theoretical diffraction-limited resolution of *λ*/NA = 670 nm. However, the sensor’s 1.85 μm pixel size imposes a practical resolution limit over five times coarser than this theoretical value, potentially introducing undersampling artifacts. To overcome this limitation, the reconstruction grid is defined with finer pixel spacing than the sensor’s pixel pitch by the factor *σ*. Paired with the operator *C*_*σ*_ {·}in the forward model [Eq. (1)], which models pixel binning at the sensor, the phase retrieval inverse problem [Eq. (2)] can achieve pixel super-resolution (PSR) for *σ* > 1, and thus reconstruct features beyond the native resolution of the sensor. This is demonstrated in **Figure 4(c)** and **4(d)**, which show reconstructions of the USAF target without and with PSR, respectively. Notably, Group 8 elements 2 (1.95 μm), 3 (1.74 μm), and 4 (1.55 μm) are only clearly resolved when PSR is applied for *σ* = 4.

### C. Calibrating spatially-varying illumination angles

A common approach in literature [30, 51] estimates LED illumination angles by cross-correlating the diffraction image from each LED with a reference image and using the resulting spatial shifts to assign fixed illumination angles, assuming ideal plane-wave illumination across the FOV. While effective for small FOVs [30, 37, 47, 49-51] (< 20 mm^2^), this assumption breaks down over larger FOVs (e.g., 40 mm^2^ in this paper), as LEDs emit spherical, not planar, wavefronts, which cause angular variation across the sample and introduces reconstruction artifacts, especially at the periphery. To address this, we introduce a spatially adaptive calibration strategy that divides the FOV into smaller subregions, each approximated as receiving local plane-wave illumination. Within each subregion, independent angle estimation and phase retrieval are performed as shown in **Figure 5**. This localized approach compensates for the spatial variation in illumination, significantly reducing reconstruction artifacts and enabling consistent image quality across large FOVs.

**Figure 5:**
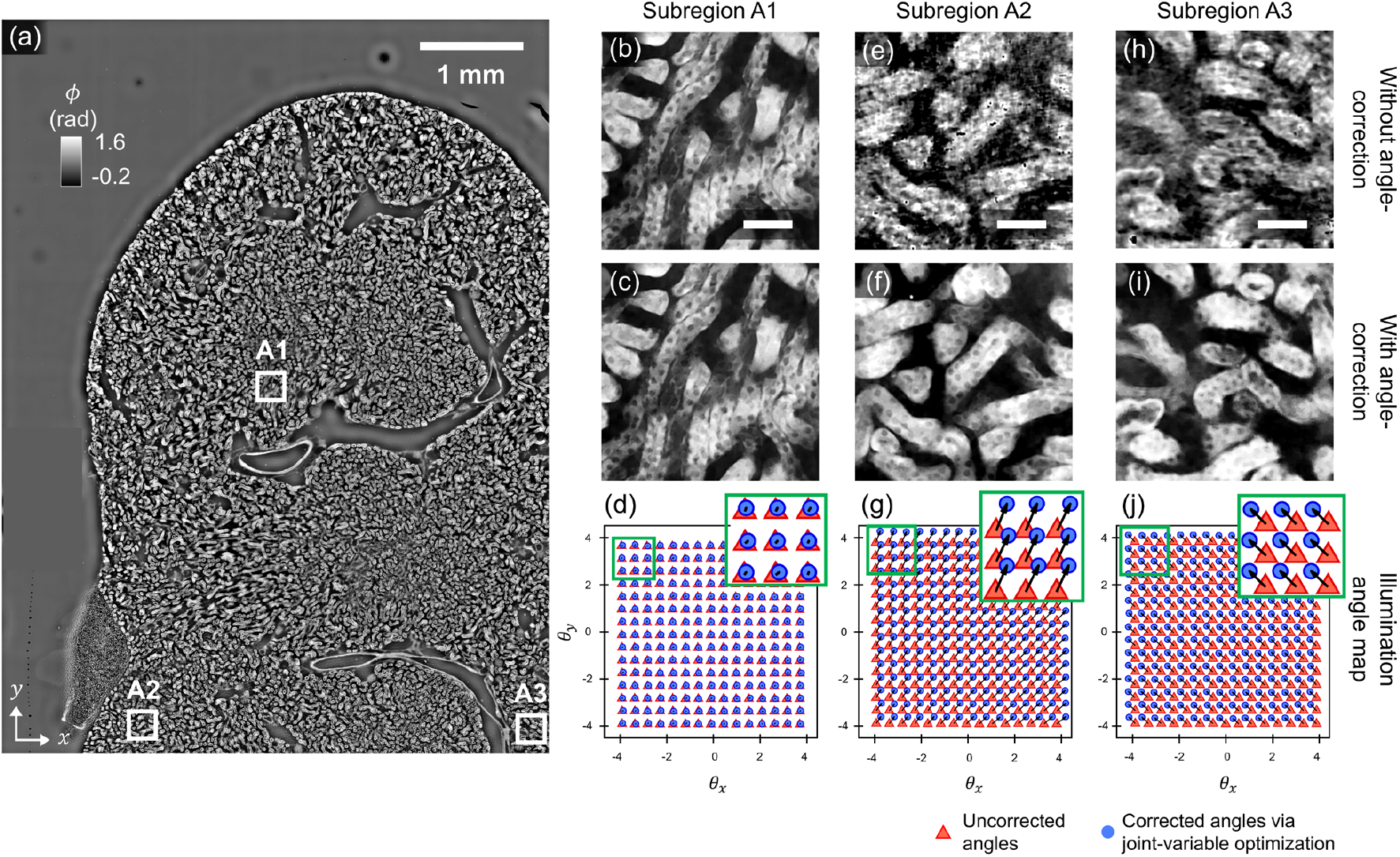
Spatially varying angle correction improves large-area phase imaging. **(a)** Full-field phase image of a 16 µm thick mouse kidney section with three subregions marked. **(b–d)** Center subregion (**A1**): reconstructions without **(b)** and with **(c)** angle correction are similar; angle map in **(d)** shows minimal variation. **(e–g)** Peripheral subregion (**A2**): correction removes artifacts (**f** vs. **e**); angle map **(g)** shows clear variation. **(h–j)** Similar improvement in another peripheral subregion (**A3**). Red triangles: initial angle estimates; blue circles: refined angles. Scale bar in subregions: 50 µm.

We validate this method using a dense 16 μm-thick C57 mouse kidney section (Invitrogen) which fills the entire sensor area. **Figure 5(a)** shows the reconstructed phase image across one sensor-FOV (7.4 mm × 5.6 mm). Three subregions are highlighted: **A1** at the center, and **A2** and **A3** near the bottom-left and bottom-right edges, respectively. In **A1**, which lies near the optical axis, reconstructions with and without angle correction [**Figure 5(b,c)**] are nearly identical, and the estimated illumination angles [**Figure 5(d)**] show minimal variation. This confirms that conventional angle estimation is sufficient near the center of the LED array, where angular variation is negligible. In contrast, reconstructions in the peripheral regions (**A2** and **A3**) exhibit significant degradation without angle-calibration, as shown in **Figure 5(e,h)**. Applying spatially varying angle correction [**Figure 5(f,i)**] effectively eliminates these artifacts and dramatically improves image quality. The corresponding corrected illumination angle maps for **A2** and **A3** are shown in **Figure 5(g)** and **5(j)**, respectively, and clearly shows spatial variation. This result demonstrates that spatially varying illumination angle correction is essential for accurate, high-quality phase imaging across large FOVs.

### D. Centimeter-scale on-chip biological phase imaging

We now demonstrate our proposed framework for high-SBP biological phase-imaging across a variety of transparent, unlabeled tissue sections (prepared as described in **Supplementary Note S6**). These samples typically spanned several centimeters, exceeding the sensor’s FOV. To image the entire sample, we captured overlapping measurements by repositioning the sensor and then stitched them using a distance-based weighting approach [57]. This method blends overlapping regions by assigning higher weights to central areas where signal quality is typically better and reducing edge contributions, thereby minimizing stitching artifacts and enabling full-FOV phase reconstruction (see **Supplementary Notes S7** and **S8**). Below, we present imaging results from tissue sections of mouse kidney, zebrafish head, and a full mouse embryo. Additional results, including tissue sections from the mammalian jejunum, lung, ovary, spinal cord, stratified epithelium, liver, and cuboidal epithelium, are provided in **Supplementary Note S9, Figures S7-S13**.

#### Adult mouse kidney

We first present high-SBP imaging results from two sagittal tissue sections of a C57 mouse kidney (Invitrogen), shown in **Figure 6(a)**, visualized across a total area of ∼1.2 cm × 1.5 cm with micron-scale imaging resolution (∼1 gigapixel). Each tissue section required measurements to be merged across four overlapping sensor FOVs. Label-free contrast clearly visualizes the anatomical organization of the kidney, including the cortex, outer and inner medulla, papilla, and pelvic cavity. Magnified views of eleven selected subregions (**A1–A11**, indicated in **Figure 6(b-l)**) highlight the system’s ability to resolve fine microanatomical features. In the cortical region spanned in the top tissue section, we observe renal cells (**A1**), distal tubules (**A2**), capillaries (**A3**), loops of Henle (**A4**), proximal tubules (**A5**), and tubular lumen (**A6**) as well as papillary draining ducts (**A7**), the outer-to-inner medulla boundary (**A8**), blood vessels (**A9**), collecting ducts (**A10**), and additional distal tubules (**A11**). These results underscore the utility of this method for histology and large-scale tissue imaging, particularly in pathology applications and systems-level tissue atlasing.

**Figure 6:**
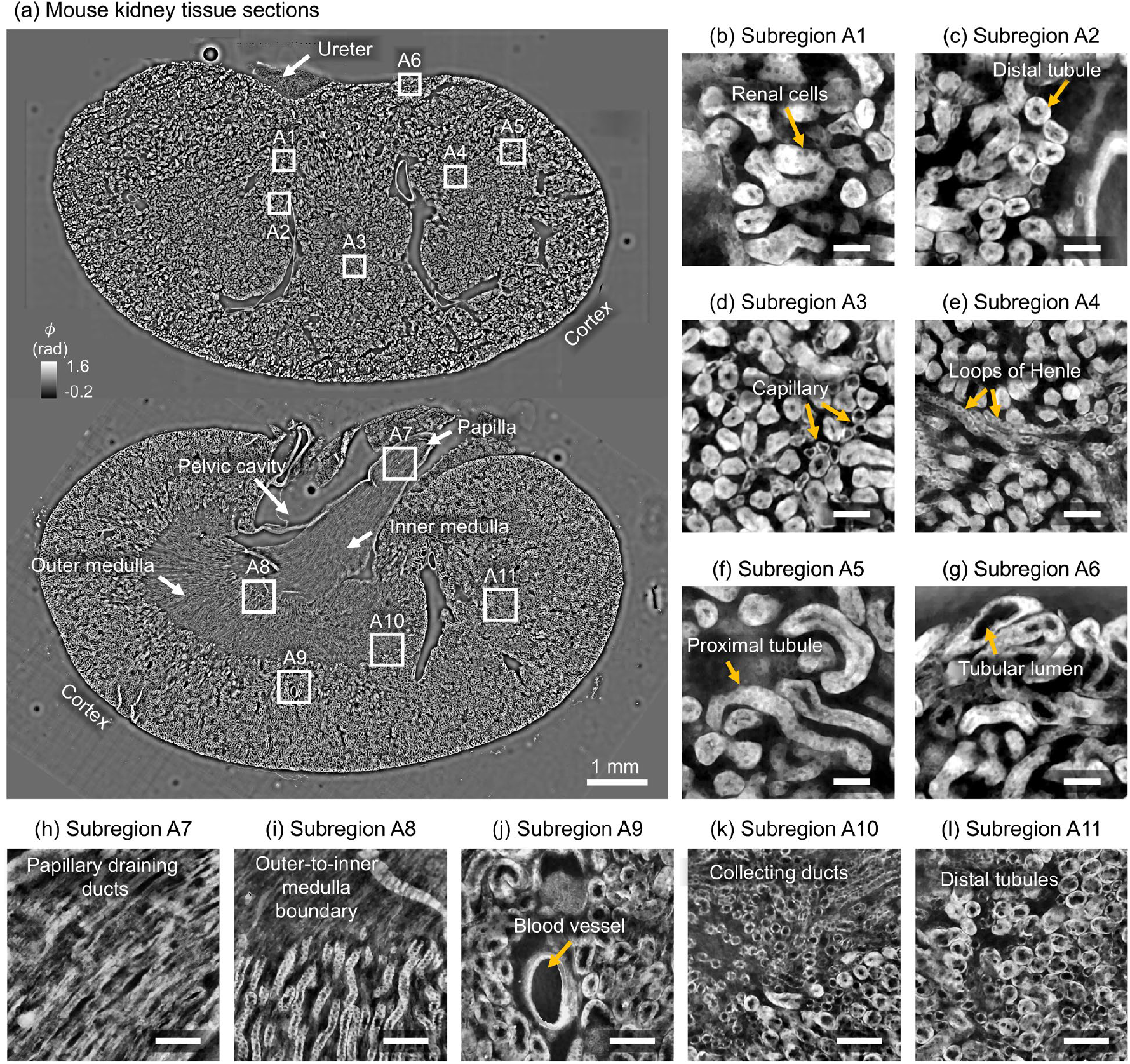
**(a)** Wide-field phase image of two sagittal kidney sections (∼1.2 cm × 1.5 cm), with subregions **A1**–**A11** highlighted. **(b–g)** Cortical subregions show renal microstructures, including renal cells (**A1**), distal tubules (**A2**), capillaries (**A3**), loops of Henle (**A4**), proximal tubules (**A5**), and tubular lumens (**A6**). **(h–l)** Medullary subregions display papillary ducts (**A7**), medullary boundaries (**A8**), blood vessels (**A9**), collecting ducts (**A10**), and distal tubules (**A11**). Quantitative phase values (φ) are shown in radians; Scale bars: 50 μm for (b–g), 100 μm for (h–l).

#### Cross-sections of larval zebrafish head

We next imaged a sample consisting of 12 μm-thick tissue sections extracted from various depths through an 5dpf (days post fertilization) zebrafish head, arranged in rows on a slide. Such samples are valuable for neuroanatomical studies due to their ability to preserve spatial organization across brain and eye structures. To image the entire specimen spanning 2.96 cm × 0.55 cm, we acquired measurements from four overlapping sensor regions and stitched them to reconstruct the full high-resolution phase image, as shown in **Figure 7(a)**.

**Figure 7:**
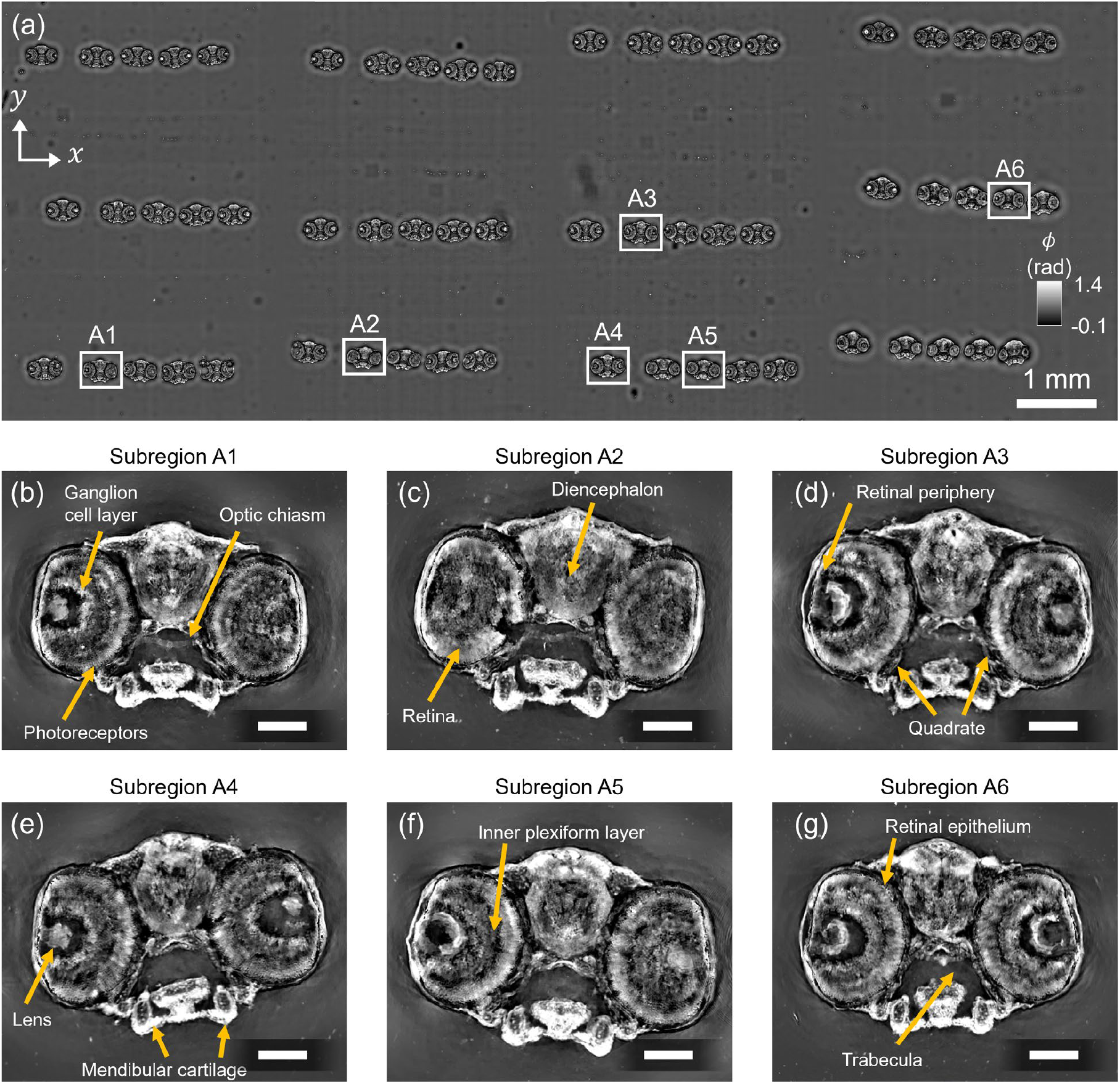
**(a)** Reconstructed high-SBP phase image of multiple transverse sections of a larval (5dpf) zebrafish head and eyes. Magnified views show detailed visualization of anatomical features such as **(b)** ganglion cell layer, photoreceptors, and the optic chiasm (**A1**); **(c)** diencephalon and retina (**A2**); **(d)** retinal periphery and quadrate (**A3**); **(e)** lens and mandibular cartilage (**A4**); **(f)** inner plexiform layer (**A5**); and **(g)** retinal pigment epithelium and trabecula (**A6**). The reconstruction highlights fine tissue features across a centimeter-scale FOV with uniform resolution. Scale bar: 100 µm.

High-resolution magnified views of selected subregions (**A1**-**A6**) in the phase reconstruction are shown in **Figure 7(b-g)**, and highlight distinct anatomical features across the brain and eye. Notable retinal structures include the ganglion cell layer, photoreceptor layer, and optic chiasm (**A1**); the inner plexiform layer (**A5**); and the retinal pigment epithelium (**A6**). Additional ocular features such as the lens (**A4**) are clearly visualized, along with brain regions like the diencephalon (**A2**) and craniofacial structures including the trabecula (**A6**), quadrate (**A3**), and mandibular cartilage (**A4**). These results demonstrate our method’s ability to maintain high image quality and resolution of structurally complex biological samples across centimeter-scale FOVs. The capacity to visualize retinal and neural microstructures without staining or labeling emphasizes this platform’s potential for ocular research, developmental biology, and neuroanatomical studies where label-free imaging would help in assessing eye screens [58] and craniofacial screens [59].

#### Mouse embryo at E18 developmental stage

Lastly, we imaged a sagittal section of a mouse embryo at embryonic day 18 (E18), a developmental stage marked by extensive anatomical complexity and advanced tissue differentiation. The tissue section, spanning approximately 2.7 cm × 1.7 cm, was reconstructed into a 2.4-gigapixel quantitative phase image by stitching 10 overlapping sensor fields of view. As shown in **Figure 8(a)**, the wide FOV reconstruction captures a diverse range of anatomical structures at micron resolution across the entire centimeter-scale embryo sample. Major regions, such as the head, thorax, abdomen, and pelvis, are clearly visualized. High-resolution zoom-ins from selected subregions [**Figure 8(b–m)**] reveal fine tissue architecture at cellular levels across various soft tissues and developing organs, including the brain, thymus, liver, midgut, and vascular structures [60]. In the cranial region, we resolve eye sockets and the olfactory lobe (**A1**) and densely packed brain cells (**A2**). In the thoracic cavity, we observe vertebral cartilage (**A3**), early bone formation within cartilage (**A4**), thymus glands and tracheal rings (**A5**), and thyroid cartilage (**A6**). Further along, we capture sternum ossification centers (**A7**), bronchioles in the developing lung (**A8**), and epithelial structures of the midgut (**A9**). In the posterior region, we visualize the vena cava (**A10**), hip joint cavity and innominate cartilage (**A11**), and the anal canal (**A12**).

**Figure 8:**
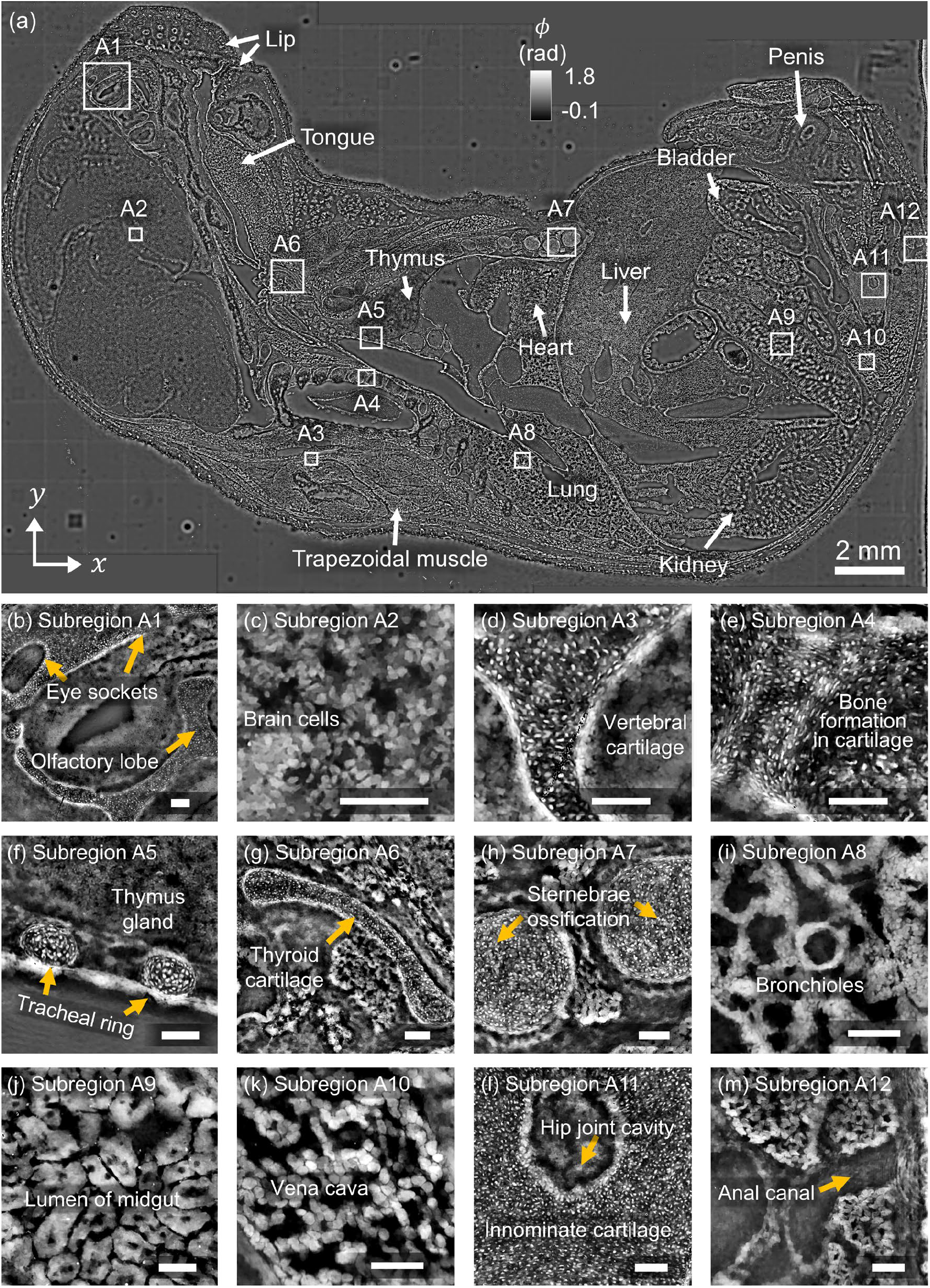
**(a)** Label-free, high-resolution phase imaging of an E18 mouse embryo sagittal section. The stitched 2.4-gigapixel image captures macroscopic structures including brain, heart, lung, liver, kidney, thymus, gut, and bladder across a 2.2 cm × 1.7 cm field. **(b–m)** Zoomed-in views show cellular and tissue-level details of soft tissues and cartilage primordia, revealing features such as tracheal rings, vertebral and thyroid cartilage, developing vasculature, and innominate cartilage at micron resolution. Scale bars: 100µm

This demonstration showcases this imaging framework’s ability to provide label-free anatomical insights across multiple biological scales, from whole-organ architecture to cellular-level detail. This capability can enable large-scale phenotyping, especially in applications involving mouse models which are often used to study genetics, development, and disease.

## 5. Discussion and conclusion

We developed a computational on-chip phase-imaging framework capable of high-resolution, large FOV imaging using angular scattering measurements with an LED array. Using this framework, we used calibration microspheres to evaluate phase accuracy, and phase resolution targets to assess imaging resolution. Our framework achieved phase accuracy with errors below 2% and delivered imaging resolution finer than the Nyquist sampling limit set by the sensor pixel size. We then conducted biological imaging experiments by demonstrating our approach on various specimens, including mouse kidney, zebrafish head, and mouse embryo tissue sections. With these demonstrations, we showed that simultaneously reconstructing the sample’s complex transmittance while also calibrating for spatially varying illumination angles for each LED source enabled ∼1 µm spatial resolution across centimeter-scale FOVs, totaling up to 2.4 gigapixels. To our knowledge, this is among the largest SBP phase-imaging demonstrations reported to date [35, 36, 61].

Looking ahead, there are several promising directions. For example, although our approach achieves spatial resolutions beyond the nominal sampling limit of the sensor pixel size, the resolution remains ultimately constrained by the pixel size rather than the optical diffraction limit. Utilizing commercially available sensors with smaller pixel sizes and larger sensor areas could help push the resolution and imaging FOV even further. Additionally, while the flat amplitude constraint effectively stabilizes the inverse problem when imaging largely-transparent samples, it does not account for amplitude variations introduced by strongly absorptive specimens. Future work could focus on developing alternative strategies to stabilize the reconstruction without relying on such a restrictive constraint. For example, this could involve collecting data under different imaging conditions, such as with sample defocus [20, 22] or structured/multiplexed illumination [50, 62]. Lastly, hardware improvements to the illumination module could further the utility of this framework for bio-imaging. For example, custom-built arrays using high-power LEDs or laser diodes could provide significantly stronger illumination. This would substantially reduce acquisition times and potentially enable live imaging applications.

Notably, our approach to 2D phase imaging relies on capturing scattering measurements of the sample being illuminated from multiple angles. Interestingly, this is precisely the same type of data used in various refractive index (RI) tomography techniques, where a sample’s 3D RI distribution is reconstructed from multi-angle measurements [57, 63, 64]. This opens up a particularly promising future direction to extend our on-chip imaging framework to 3D, enabling high-SBP and label-free volumetric imaging. In fact, our framework, as described in Eqs. (1) and (2), can be directly adapted to many iterative 3D RI solvers, where the forward model is reformulated to account for multiple scattering effects. Adapting this approach for 3D RI imaging in thicker, scattering samples would be particularly exciting, as it would allow label-free and fully volumetric visualization of complex 3D structures over ultra-large FOVs. Combined with the simplicity and cost-effectiveness of this approach, this label-free high-SBP volumetric imaging capability would broaden even further the applications of this on-chip imaging framework.

## Acknowledgment

We thank Colton Kostelnik and Manuel Rausch in the Department of Biomedical Engineering at UT Austin for assistance with preparing FFPE tissue sections. Sectioning was performed at the Center for Biomedical Research Support Microscopy and Flow Cytometry Facility at UT Austin. The Texas Advanced Computing Center (TACC) at The University of Texas at Austin provided computational resources that contributed to the research results reported within this paper. We also gratefully acknowledge Bill Shawlot at the UT Mouse Genetic Engineering Facility, Richard Levenson from UC Davis, as well as Maciej Trusiak, Mikołaj Rogalski, Piotr Arcab, and Emilia Wdowiak from the Warsaw University of Technology for insightful discussions regarding sample preparation methods and lens-free phase imaging. Additionally, we thank Diptodip Deb and Gert-Jan Both at the Janelia Research Campus for valuable discussions regarding the JAX implementation for high-speed compute.

## Data availability

Data underlying the results presented in this paper may be obtained from the authors upon reasonable request.

## Supplemental document

See Supplement 1 for supporting content.

## References

[1] D. Dan, “Frontiers | Editorial: Optical microscopy: advances and applications,” Frontiers in Physics, vol. 11, 2023/11/27, doi: 10.3389/fphy.2023.1337300.

[2] H. Balasubramanian et al., “Imagining the future of optical microscopy: everything, everywhere, all at once,” Communications Biology 2023 6:1, vol. 6, no. 1, 2023-10-28, doi: 10.1038/s42003-023-05468-9.

[3] X. Chen, B. Zheng, and H. Liu, “Optical and digital microscopic imaging techniques and applications in pathology,” Analytical Cellular Pathology (Amsterdam), vol. 34, no. 1-2, 2011, doi: 10.3233/ACP-2011-0006.

[4] C. Ferreira et al., “Space–bandwidth product of optical signals and systems,” JOSA A, Vol. 13, Issue 3, pp. 470–473, vol. 13, no. 3, 1996-03-01, doi: 10.1364/JOSAA.13.000470.

[5] J. Park, D. J. Brady, G. Zheng, L. Tian, and L. Gao, “Review of bio-optical imaging systems with a high space-bandwidth product,” Advanced photonics, vol. 3, no. 4, 2021 Jun 26, doi: 10.1117/1.ap.3.4.044001.

[6] S. A. Alexandrov et al., “High-resolution, wide-field object reconstruction with synthetic aperture Fourier holographic optical microscopy,” Optics Express, Vol. 17, Issue 10, pp. 7873–7892, vol. 17, no. 10, 2009-05-11, doi: 10.1364/OE.17.007873.

[7] G. Zheng, R. Horstmeyer, C. Yang, G. Zheng, R. Horstmeyer, and C. Yang, “Wide-field, high-resolution Fourier ptychographic microscopy,” Nature Photonics 2013 7:9, vol. 7, no. 9, 2013-07-28, doi: 10.1038/nphoton.2013.187.

[8] S. C. Shanmugavel, Y. Zhu, S. C. Shanmugavel, and Y. Zhu, “Structured illumination contrast transfer function for high resolution quantitative phase imaging,” Optics Express, Vol. 31, Issue 24, pp. 40151–40165, vol. 31, no. 24, 2023-11-20, doi: 10.1364/OE.504961.

[9] L. Waller, S. Chowdhury, L.-H. Yeh, L.-H. Yeh, S. Chowdhury, and L. Waller, “Computational structured illumination for high-content fluorescence and phase microscopy,” Biomedical Optics Express, Vol. 10, Issue 4, pp. 1978–1998, vol. 10, no. 4, 2019-04-01, doi: 10.1364/BOE.10.001978.

[10] L. Waller et al., “Speckle-structured illumination for 3D phase and fluorescence computational microscopy,” Biomedical Optics Express, Vol. 10, Issue 7, pp. 3635–3653, vol. 10, no. 7, 2019-07-01, doi: 10.1364/BOE.10.003635.

[11] “Gigapixel imaging with a novel multi-camera array microscope,” 2022-12-14, doi: 10.7554/eLife.74988.

[12] K. C. Zhou et al., “Parallelized computational 3D video microscopy of freely moving organisms at multiple gigapixels per second,” Nature Photonics 2023 17:5, vol. 17, no. 5, 2023-03-20, doi: 10.1038/s41566-023-01171-7.

[13] J. Fan et al., “Video-rate imaging of biological dynamics at centimetre scale and micrometre resolution,” Nature Photonics 2019 13:11, vol. 13, no. 11, 2019-07-08, doi: 10.1038/s41566-019-0474-7.

[14] A. Greenbaum et al., “Imaging without lenses: achievements and remaining challenges of wide-field on-chip microscopy,” Nature Methods 2012 9:9, vol. 9, no. 9, 2012-08-30, doi: 10.1038/nmeth.2114.

[15] J. Zhang, J. Sun, Q. Chen, and C. Zuo, “Resolution Analysis in a Lens-Free On-Chip Digital Holographic Microscope | IEEE Journals & Magazine | IEEE Xplore,” IEEE Transactions on Computational Imaging, vol. 6, 2020, doi: 10.1109/TCI.2020.2964247.

[16] W. Bishara et al., “Lensfree on-chip microscopy over a wide field-of-view using pixel superresolution,” Optics Express, Vol. 18, Issue 11, pp. 11181–11191, vol. 18, no. 11, 2010-05-24, doi: 10.1364/OE.18.011181.

[17] W. Luo et al., “Pixel super-resolution using wavelength scanning,” Light: Science & Applications 2016 5:4, vol. 5, no. 4, 2015-12-14, doi: 10.1038/lsa.2016.60.

[18] X. Wu et al., “Wavelength-scanning lensfree on-chip microscopy for wide-field pixel-super-resolved quantitative phase imaging,” Optics Letters, Vol. 46, Issue 9, pp. 2023–2026, vol. 46, no. 9, 2021-05-01, doi: 10.1364/OL.421869.

[19] M. Guizar-Sicairos, J. R. Fienup, M. Guizar-Sicairos, and J. R. Fienup, “Phase retrieval with transverse translation diversity: a nonlinear optimization approach,” Optics Express, Vol. 16, Issue 10, pp. 7264–7278, vol. 16, no. 10, 2008-05-12, doi: 10.1364/OE.16.007264.

[20] K. Niedziela, M. Rogalski, P. Arcab, J. Winnik, P. Zdańkowski, and M. Trusiak, “Pixel Super Resolution with Axial Scanning in Lensless Digital In-line Holographic Microscopy,” Photonics Letters of Poland, vol. 16, no. 4, 2024/12/31, doi: 10.4302/plp.v16i4.1305.

[21] Y. Chen et al., “Pixel-super-resolved lens-free quantitative phase microscopy with partially coherent illumination,” npj Nanophotonics 2024 1:1, vol. 1, no. 1, 2024-06-28, doi: 10.1038/s44310-024-00015-8.

[22] A. Ozcan, A. Greenbaum, A. Greenbaum, and A. Ozcan, “Maskless imaging of dense samples using pixel super-resolution based multi-height lensfree on-chip microscopy,” Optics Express, Vol. 20, Issue 3, pp. 3129–3143, vol. 20, no. 3, 2012-01-30, doi: 10.1364/OE.20.003129.

[23] E. Wdowiak et al., “Quantitative phase imaging verification in large field-of-view lensless holographic microscopy via two-photon 3D printing,” Scientific Reports 2024 14:1, vol. 14, no. 1, 2024-10-09, doi: 10.1038/s41598-024-74866-8.

[24] J. K. Adams et al., “In vivo lensless microscopy via a phase mask generating diffraction patterns with high-contrast contours,” Nature Biomedical Engineering 2022 6:5, vol. 6, no. 5, 2022-03-07, doi: 10.1038/s41551-022-00851-z.

[25] A. F. Coskun, T.-W. Su, I. Sencan, and A. Ozcan, “Lensless Fluorescent Microscopy on a Chip,” Journal of Visualized Experiments : JoVE, no. 54, 2011 Aug 17, doi: 10.3791/3181.

[26] J. K. Adams et al., “Single-frame 3D fluorescence microscopy with ultraminiature lensless FlatScope,” Science advances, vol. 3, no. 12, p. e1701548, 2017.

[27] G. N. McKay et al., “Lens Free Holographic Imaging for Urinary Tract Infection Screening | IEEE Journals & Magazine | IEEE Xplore,” IEEE Transactions on Biomedical Engineering, vol. 70, no. 3, 2023, doi: 10.1109/TBME.2022.3208220.

[28] S. Jiang et al., “Wide-field, high-resolution lensless on-chip microscopy via near-field blind ptychographic modulation,” Lab on a Chip, vol. 20, no. 6, 2020/03/17, doi: 10.1039/C9LC01027K.

[29] Y. Fang et al., “High-Precision Lens-Less Flow Cytometer on a Chip,” Micromachines 2018, Vol. 9, Page 227, vol. 9, no. 5, 2018-05-10, doi: 10.3390/mi9050227.

[30] W. Luo et al., “Synthetic aperture-based on-chip microscopy,” Light: Science & Applications 2015 4:3, vol. 4, no. 3, 2015-03-27, doi: 10.1038/lsa.2015.34.

[31] A. Greenbaum et al., “Wide-field computational imaging of pathology slides using lens-free on-chip microscopy,” Science Translational Medicine, vol. 6, no. 267, 2014-12-17, doi: 10.1126/scitranslmed.3009850.

[32] Y.-C. Wu et al., “Air quality monitoring using mobile microscopy and machine learning,” Light: Science & Applications 2017 6:9, vol. 6, no. 9, 2017-03-15, doi: 10.1038/lsa.2017.46.

[33] J. Zhang et al., “Adaptive pixel-super-resolved lensfree in-line digital holography for wide-field onchip microscopy,” Scientific Reports 2017 7:1, vol. 7, no. 1, 2017-09-18, doi: 10.1038/s41598-017-11715-x.

[34] C. Xu, A. Cao, H. Pang, Q. Deng, S. Hu, and H. Yang, “Lensless imaging via multi-height mask modulation and ptychographical phase retrieval,” Optics and Lasers in Engineering, vol. 169, 2023/10/01, doi: 10.1016/j.optlaseng.2023.107739.

[35] S. Jiang et al., “Resolution-Enhanced Parallel Coded Ptychography for High-Throughput Optical Imaging,” ACS Photonics, vol. 8, no. 11, October 14, 2021, doi: 10.1021/acsphotonics.1c01085.

[36] S. Jiang et al., “Blood-Coated Sensor for High-Throughput Ptychographic Cytometry on a Blu-ray Disc,” ACS Sensors, vol. 7, no. 4, April 8, 2022, doi: 10.1021/acssensors.1c02704.

[37] L. Cao, Y. Gao, Y. Gao, and L. Cao, “Generalized optimization framework for pixel super-resolution imaging in digital holography,” Optics Express, Vol. 29, Issue 18, pp. 28805–28823, vol. 29, no. 18, 2021-08-30, doi: 10.1364/OE.434449.

[38] P. Gao, G. Pedrini, and W. Osten, “Structured illumination for resolution enhancement and autofocusing in digital holographic microscopy.,” Optics Letters, vol. 38, no. 8, 2013, doi: 10.1364/OL.38.001328.

[39] J. Bae et al., “Design and single-shot fabrication of lensless cameras with arbitrary point spread functions,” Optica, Vol. 10, Issue 1, pp. 72–80, vol. 10, no. 1, 2023-01-20, doi: 10.1364/OPTICA.466072.

[40] F. Zhang et al., “Phase retrieval using multiple illumination wavelengths,” Optics Letters, Vol. 33, Issue 4, pp. 309–311, vol. 33, no. 4, 2008-02-15, doi: 10.1364/OL.33.000309.

[41] C. Zuo et al., “Lensless phase microscopy and diffraction tomography with multi-angle and multiwavelength illuminations using a LED matrix,” Optics Express, Vol. 23, Issue 11, pp. 14314–14328, vol. 23, no. 11, 2015-06-01, doi: 10.1364/OE.23.014314.

[42] J. A. Picazo-Bueno et al., “Improved quantitative phase imaging in lensless microscopy by singleshot multi-wavelength illumination using a fast convergence algorithm,” Optics Express, Vol. 23, Issue 16, pp. 21352–21365, vol. 23, no. 16, 2015-08-10, doi: 10.1364/OE.23.021352.

[43] L. Waller, L. Tian, L. Tian, and L. Waller, “3D intensity and phase imaging from light field measurements in an LED array microscope,” Optica, Vol. 2, Issue 2, pp. 104–111, vol. 2, no. 2, 2015-02-20, doi: 10.1364/OPTICA.2.000104.

[44] K. Rasul et al., “Lensless on-chip LED array microscope using amplitude and phase masks,” JOSA B, Vol. 37, Issue 12, pp. 3652–3659, vol. 37, no. 12, 2020-12-01, doi: 10.1364/JOSAB.396076.

[45] J. R. Fienup, C. C. Wackerman, J. R. Fienup, and C. C. Wackerman, “Phase-retrieval stagnation problems and solutions,” JOSA A, Vol. 3, Issue 11, pp. 1897–1907, vol. 3, no. 11, 1986-11-01, doi: 10.1364/JOSAA.3.001897.

[46] P. Arcab, M. Rogalski, and M. Trusiak, “Single-shot experimental-numerical twin-image removal in lensless digital holographic microscopy,” Optics and Lasers in Engineering, vol. 172, 2024/01/01, doi: 10.1016/j.optlaseng.2023.107878.

[47] Z. Zhang et al., “Invited Article: Mask-modulated lensless imaging with multi-angle illuminations,” APL Photonics, vol. 3, no. 6, 2018/06/01, doi: 10.1063/1.5026226.

[48] J. Kim et al., “Ptychographic lens-less birefringence microscopy using a mask-modulated polarization image sensor,” Scientific Reports 2023 13:1, vol. 13, no. 1, 2023-11-07, doi: 10.1038/s41598-023-46496-z.

[49] S. Jiang et al., “Mask-modulated lensless imaging via translated structured illumination,” Optics Express, Vol. 29, Issue 8, pp. 12491–12501, vol. 29, no. 8, 2021-04-12, doi: 10.1364/OE.421228.

[50] G. Zheng et al., “Single-shot lensless imaging via simultaneous multi-angle LED illumination,” Optics Express, Vol. 26, Issue 17, pp. 21418–21432, vol. 26, no. 17, 2018-08-20, doi: 10.1364/OE.26.021418.

[51] P. Li, A. Maiden, P. Li, and A. Maiden, “Lensless LED matrix ptychographic microscope: problems and solutions,” Applied Optics, Vol. 57, Issue 8, pp. 1800–1806, vol. 57, no. 8, 2018-03-10, doi: 10.1364/AO.57.001800.

[52] E. L. Dixon et al., “Simple and robust image-based autofocusing for digital microscopy,” Optics Express, Vol. 16, Issue 12, pp. 8670–8677, vol. 16, no. 12, 2008-06-09, doi: 10.1364/OE.16.008670.

[53] J. W. Goodman, Introduction to Fourier Optics. Greenwood village, CO: Roberts & Company Publishers, 2005.

[54] S. T. Thurman, M. Guizar-Sicairos, J. R. Fienup, M. Guizar-Sicairos, S. T. Thurman, and J. R. Fienup, “Efficient subpixel image registration algorithms,” Optics Letters, Vol. 33, Issue 2, pp. 156–158, vol. 33, no. 2, 2008-01-15, doi: 10.1364/OL.33.000156.

[55] V. Elser and V. Elser, “Phase retrieval by iterated projections,” JOSA A, Vol. 20, Issue 1, pp. 40–55, vol. 20, no. 1, 2003-01-01, doi: 10.1364/JOSAA.20.000040.

[56] D. Johnson, P. Li, A. Maiden, A. Maiden, D. Johnson, and P. Li, “Further improvements to the ptychographical iterative engine,” Optica, Vol. 4, Issue 7, pp. 736–745, vol. 4, no. 7, 2017-07-20, doi: 10.1364/OPTICA.4.000736.

[57] R. Eckert et al., “High-resolution 3D refractive index microscopy of multiple-scattering samples from intensity images,” Optica, Vol. 6, Issue 9, pp. 1211–1219, vol. 6, no. 9, 2019-09-20, doi: 10.1364/OPTICA.6.001211.

[58] J. Lee et al., “An ENU Mutagenesis Screen in Zebrafish for Visual System Mutants Identifies a Novel Splice-Acceptor Site Mutation in patched2 that Results in Colobomas,” Investigative Ophthalmology & Visual Science, vol. 53, no. 13, 2012 Dec 13, doi: 10.1167/iovs.12-11061.

[59] N. SC et al., “Mutations affecting craniofacial development in zebrafish - PubMed,” Development (Cambridge, England), vol. 123, no. 1, 1996 Dec, doi: 10.1242/dev.123.1.357.

[60] M. H. Kaufman, The Atlas of Mouse Development, 2 ed. San Diego: Academic Press, 1992.

[61] J. Park et al., “Review of bio-optical imaging systems with a high space-bandwidth product,” Advanced Photonics, vol. 3, no. 4, 2021/06, doi: 10.1117/1.AP.3.4.044001.

[62] S. Jiang et al., “Mask-modulated lensless imaging via translated structured illumination,” Optics Express, Vol. 29, Issue 8, pp. 12491–12501, vol. 29, no. 8, 2021-04-12, doi: 10.1364/OE.421228.

[63] J. Kim et al., “DMD and microlens array as a switchable module for illumination angle scanning in optical diffraction tomography,” Biomedical Optics Express, Vol. 15, Issue 10, pp. 5932–5946, vol. 15, no. 10, 2024-10-01, doi: 10.1364/BOE.535123.

[64] J. Kim, S. Chowdhury, J. Kim, and S. Chowdhury, “Space-time inverse-scattering of translation-based motion,” Optica, Vol. 12, Issue 5, pp. 643–653, vol. 12, no. 5, 2025-05-20, doi: 10.1364/OPTICA.554264.

